# Unraveling Evolutionary Dynamics: Insights from In Silico Experiments on Selective Mechanisms in Controlled Environments

**DOI:** 10.1101/2023.10.24.563778

**Authors:** Marco Ledda, Alessandro Pluchino, Marco Ragusa

**Author notes:** Contributing authors.

## Abstract

In this paper, we present a series of *in silico* experiments aimed at probing the evolutionary properties of our model. Our investigation encompasses multiple methodologies, beginning with the standard model used in population genetics for measuring natural selection. Next, we employ the Price equation, a well-established formalism known for its effectiveness in tracking how the relationship between parents and offspring evolves over time. In conclusion, we delve into the model results to explain, in the light of evolutionary theory, how the selective mechanism operates. Furthermore, the speculation about the mechanism will be hindered on the agents of the selective process. Even though natural selection can be considered as a statistical phenomenon, sprouting from the change in population frequencies, we argue that in models where there is a elevate control on the environment, it is possible to define the single element responsible for the selective pressure on the *units of selection*.

## 1 Introduction

In modern scientific research, the primary goal is to uncover the dynamic laws governing the behavior of various systems, whether they are physical, chemical, biological, or social. These laws serve as the Newtonian mathematical framework for efficient causation, one of the four Aristotelian causes which account for the mechanism of the occurrence of some, relatively simple, phenomena^1^[1]. Complex systems, on the other hand, present a distinct challenge. They comprise a multitude of entities from various hierarchical levels, engaging in intricate interactions that defy a straightforward explanation through efficient causation. This complexity is particularly pronounced in biological systems, where entities span different levels, with properties and dynamical relations shifting as one moves from one level to another [2]. Consider, for example, the relationship between structure and function, which underscores that comprehending, or replicating, a particular function from the corresponding structure — be it network topology, biochemical attributes, or three-dimensional shape — does not elucidate how the same structure can perform new functions in disparate environments. Such peculiarities entail the appeal to a more sophisticated account of causation in complex systems, or, at least, an updated version of efficient causation, [3].

Biocomplexity research has highlighted how simple local rules can engender complex phenomena at higher organizational levels. For instance, the movement of a single cell can be influenced by binary or gradient signals, gene transcription lies on the presence or absence of specific transcription factors, and elementary foraging behaviors can lead to intricate trophic networks in ecological systems. Living organisms, in essence, constitute dynamic agglomerations of matter, energy, and information. To delve into the manner in which organisms organize matter and exploit energy by deciphering information, is an endeavor that involves dissecting these entities into their constituent components, studying their unique attributes, and subsequently reconstructing their interrelations, [4, 5]. A powerful way to comprehend biological systems is to reproduce them, or at least some of their characteristics, with computational methods, where relevant variables are already chosen, and environmental fluctuations are more easily manageable. However, conducting experiments in fields deeply rooted in history has always been a challenging task. In classical experimental sciences like physics, chemistry, or molecular biology, scientists work with a limited number of variables in a controlled environment, assuming that both endogenous and exogenous perturbations of the system can be integrated into the analysis. Closed environments like labs or individual experiments allow for this level of control and the repeated testing of the same process while modifying one or more variables each time. However, when a historical dimension is inherent to the system, a significant challenge arises: it becomes impossible to intervene multiple times without altering the evolutionary history of the entity in question, [6].

Biological systems often span over extended time scales, with varying rates of change, such as those observed in macroevolution and microevolution. Moreover, once a specific system’s evolutionary history has unfolded, it becomes impossible to restart the process by altering one or more variables in order to acquire causal insights, infact, attempting to replay the historical process would yield different outcomes, as discussed by [7]. One particular category of computational models, specifically agent-based models, offers a valuable approach for examining such processes. These models enable the compression of time scales and the selection of relevant variables, rendering them more manageable for researchers. Additionally, they facilitate the exploration of spatial dynamics, which are essential for understanding evolutionary changes in ecological and population-based contexts.

The main objective of this study is to utilize agent-based modeling to simulate the behavior of a specific biological system, colorectal cancer, [8]. Our aim is to investigate the evolutionary characteristics of the system and how they respond to interventions. The following sections will cover: evolutionary methods in cancer research, an analysis of the model using standard population genetics and Price’s equation, and a brief discussion about the generalization of model findings and the mechanism of natural selection.

## 2 Evolutionary methods in cancer research

Since the mid-1970s, cancer development has been understood as a Darwinian process of mutation, selection, and adaptation, in which the most proliferative and uncontrolled cells lead to the development of neoplastic tissue [9]. The simplest explanation for this phenomenon is that a random mutation creates a cell with a proliferation rate advantageous compared to the other cells. This cell and its descendants then proliferate more efficiently than their neighbors, eventually becoming a large clonal population that dominates the tissue [10]. However, this process is an oversimplification that fails to capture the complexity of cancer development. Mutations can occur not only in a single cell, but also in different cells at different moments during the organism’s life, resulting in multiclonal population expansion rather than a single clonal one. This means that cancer is not just a monoclonal disease, but rather a heterogeneous mixture of different cell populations with distinct genetic and epigenetic profiles [11, 12]. Despite these challenges, evolutionary theory has played an ever-growing role in analyzing cancer development and progression, helping researchers to understand the dynamic interplay between genetic and environmental factors that shape the evolution of cancer cell populations [13, 14]. Although the fundamental process is constitute by somatic mutations and evolution from the wild-type class of alleles, eventually leading to metastasis and resistance to therapies, there is a profound lack of knowledge about how the malignacy is reached in the different tumors and patients, [15]. Some hypothesis pointed out the extreme plasticity in adaptation to new environments, or the capacity to assume stem cells-like characteristics, which means to have a powerful functional modularity in face of the environment, [16]. Over the last two decades we have assisted a flourishing development of evolutionary and ecological studies geared to the the analysis of tumor development, the discovery of new therapeutic treatment, and the empowerment of clinical control [13, 17, 18]. Many studies, conducted following evolutionary theory methods, helped cancer research to estimate crucial parameters, both evolutionary and ecological: birth and death rates, driver mutation rate, drug induced death rate, mutation rate, population size, hypoxic niche, migration rates, vascularization of the tumoral agglomerate, [19]. Among the various mechanisms that drive evolution in biological systems, selection, mutation, random drift, to name a few, there is debate about the centrality of selective process in the take over of tumoral *renegade* cells, and the pervasiveness of a neutral process, whom view is that neutral molecular evolution is the major, not the unique, factor of differentiation among cellular neoplastic populations, [20]. However, it is true that while the vast part of molecular variation is neutral respect of selective pressure, driver mutations give fitness advantage to clones and sub-clone who bear the same genotype, and fitness difference is the hallmark of selective mechanism.

Evolution involves inheritance of traits with variation, and in the organism this evolution is observable at the cellular level, either individual or of the population. Differences in individuals (cells) lead to the phenomenon known as *ITH-intratumoral heterogeneity*, derived by the accumulation of mutation in the DNA, due to alterations in environmental conditions (i.e., hipoxya, inflammation, etc.), and in the epigenetic package, in oter words, to genetic instability factors, [14, 21]. Therefore, evolutionary theory can play a relevant role in the detection of lineage relations, selective pressure, or neutral processes, which leads to a proper speciation process within the tissue and outside it when metastasis its involved, [22]. Moreover, radically important in the treatment of a given tumor in a specific patient, is the field of precision medicine, focused on the discovery of the ways to predict how the single-patient genomic line would give information about, for example, which drugs would target the specific tumor or if the cancer will comes back after the therapy, [23].

Mathematical models are powerful tools aimed to find the dynamical invariants within a system and between system components. In cancer research those methods have been extremely useful to trace population dynamic, phylogenetic trees inference, and fitness landscapes of discovered driver genes, [24]. Some works on CRC analysis have shown the relevance of this approach to account for genetic and environmental features of this specific cancer. In [25], authors delve into the genome analysis of CRC finding that, remarkably, the majority of intratumoral variation is due to the plasticity of cells in function of their environment, rather than for the contribution of an ancestral mutation, reverberated in the sub-clonal population. The distinction between driver mutation and neutral or hitchhiking mutations is another central problem of cancer studies, and authors in [26], tried to replicate the mutational accumulation process simulating a two phase model. In the first phase the population is in a homeostatic state with cells are proliferating and are subjected to a probability of mutation. Once the number of additive mutation overtake a given threshold, the second sub-clonal expansion phase begins. The final result has been the design of a hitchhiking index to measure the probability of a gene to be a passenger mutation rather then a driver one. As said before, evolutionary analysis of cancer is useful also to shed light on the environmental features of neoplastic phenomena. In [27] authors pointed out that phenotypic malignant traits arise in the context of an altered microenvironment, in particular that adaptation to hipoxya and augmented acidosis, is the special factor yielding to the last stages of carcinogenesis.

## 3 Detecting selection

### 3.1 Classic Population Genetics

Population genetics is a field that revolves around tracking changes in gene frequency over successive generations and involves modeling various fundamental mechanisms such as natural selection, random genetic drift, and migration. It is noteworthy that alterations in gene frequencies can arise from gradual mutations in genes that are either neutral or nearly neutral, considering only a small fraction of the entire genetic material as adaptive or responsive to adaptation, as discussed by Kimura (1989) and Kimura (1983) [28, 29].

However, for the purposes of this study, our focus will be exclusively on natural selection, as it stands out as the most pertinent mechanism when we are addressing adaptation to novel environmental conditions and fitness differences among individuals. To model genetic variation within a population, the simplest approach entails considering a single genetic locus with two allelic forms, denoted as *A*_1_ and *A*_2_, from which three diploid genotypes arise: *A*_1_, *A*_1_; *A*_1_, *A*_2_; and *A*_2_, *A*_2_. Each genotype is assigned a fitness value, represented as *w*_11_, *w*_12_, and *w*_22_ which quantifies the average number of successful gametes contributed to future generations by each specific genotype. The formalization of these idealized genetic entities is due to G.H. Hardy and W. Weinberg whose contributions has been to assign algebraic values to the allelic frequencies, in the following way

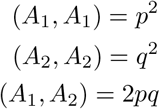

Hardy and Weinberg proposed the concept of the Hardy-Weinberg equilibrium, which occurs when a population’s allelic frequencies remain stable over generations, provided that certain key conditions are met. This equilibrium holds true only if all the following criteria are satisfied:

**No Selection**: This implies that no individual within the population has a differential survival or reproductive advantage over others. In other words, there is no natural selection favoring one genotype over another.

**No Mutations**: There are no new variations or mutations arising from the wild-type allele. Allelic frequencies remain constant, and there is no introduction of novel genetic variants.

**No Migration**: The population remains closed, with no influx or efflux of individuals from other populations. It remains isolated, and genetic exchange with other groups does not occur.

**Random Mating**: In sexually reproducing populations, individuals exhibit no preference in selecting their reproductive partners. Mate choice is entirely random.

When any one of these assumptions is violated, the population deviates from the equilibrium, and this deviation is particularly relevant when examining the extent of variation between generations due to natural selection. To quantify this variation accurately, it is necessary to employ the appropriate formalism or mathematical framework.

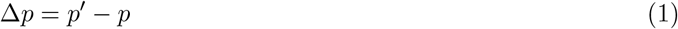

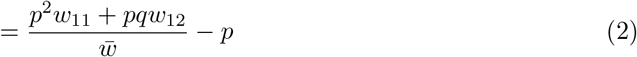

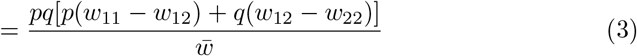

This is the fundamental model of Natural Selection, and when this quantity is represented in function of the wild-type allele *p*, what is observed is how strong or weak is the variation for each frequency of the allele. If Δ*P >* 0 then some natural selection has led *A*_1_ to increase its frequency in the population, conversely, if Δ*P <* 0 then the selection process has led *A*_1_ to a decrease in the population, finally, if Δ*P* = 0 there is no change in the allelic frequency, [30].

Even if in classic population genetics, it is common to assume that populations exhibit sexual reproduction with random mating, feature which is not present in our model, we can argue that this case can be effectively treated as if it incorporates these characteristics. To address the assumption of random mating, which ensures no preferential mating and no preferential direction in allele competition, we can equate this requirement with the hypothesis that cells in our model mate only with individuals bearing the same genotype. This is always true since cells essentially reproduce copies of themselves. Regarding the recombination factor, which implies that only complete deletions or point mutations are allowed, this is readily achievable too. The individuals (cells) undergo mitotic activities, giving rise to clones, without any genetic exchange interaction with other cells. They are subjected to random mutations, but the genetic material remains intact without recombination. The only challenge arises from the assumption of non-overlapping populations, which our model does not fully satisfy because cells that enter the mitotic phase do not die after the first mitotic activity. Consequently, there will always be a proportion constituted not just by the contribution of *p* to the next generation but also by *p* itself. In a model where stochastic elements play a crucial role, it would be unwise to disregard the possibility that certain allelic variants may become predominant purely by chance. Genetic drift is a neutral effect, meaning that there is no preferential direction in trait fixation. Depending on the population size, it takes a certain amount of time for a particular allele to fixate in the population [31]. While the Wright-Fisher model is commonly used to calculate genetic drift, representing the transition from one generation to another with a binomial distribution [32], we will employ some derivations of the Moran model. The Moran model is a stochastic process used to track the evolution of a system with two or more types of reproducing individuals, subject to mutational events, and it allows for population overlapping [33].

Consider a constant population in the crypt, denoted as N = 2500, with two types of individuals: physiological *a* and a mutant variant *b*. The mutation rate, set to *u* = 10^*−*9^, implies that the probability of reproduction for a individuals without mutations is 1*−u*, while the reproduction rate for b phenotypes is *Nu*. Let’s assume that *b* has a relative fitness *r* compared to the fitness value of a individuals, which is represented as 1*/r*. In our model, the fitness value of physiological cells is 1/24 hours, and the first mutated phenotype that changes cellular fitness is *adenoma* with a fitness of 1/12 hours. Following from these assumptions, the probability that *b* will outcompete a can be calculated as:

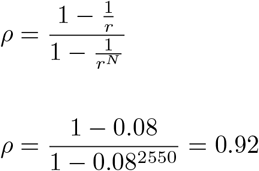

Once we have the transition probability from a state to another, we can calculate the rate of evolution from *a* to *b*, which is *R* = *Nuρ* = 0.0000023, thus, a really low rate of transition. If *b* is neutral, i.e., does not affect the fitness value, then the rate is equals to *u*. The key conclusion is that when there is a fitness difference, the population element can indeed play a role in reducing the rate at which new variants take over. However, a more crucial one, is the probability of mutation that, when is high, augment the likelihood of an increased frequency of *b*, whilst, when is at a physiological value is not sufficient for the fixation, even in the mean lifespan of a human being

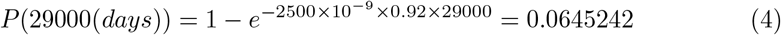

Furthermore, if the introduced mutations are entirely neutral, the rate is solely determined by the probability of mutation, which in the physiological scenario is extremely low. These calculations prompt speculation about the environmental and epigenetic factors that contribute to an increased probability of mutation, in fact, these results assume greatly higher probabilities of occurrence with the increase the term *u*. In reality, this phenomenon arises from a complex interplay of various factors, which is a feature not modeled in this work, except for the mutation of *P53* which, when mutated, augment the probability of mutation in the other two genes, [34–36].

### 3.2 Gene frequencies in the crypt

If the assumptions we’ve made so far remain valid, we can now instantiate the variables of the natural selection model with the genes involved in colorectal cancer (CRC) progression, as represented in our model of the colonic crypt. We have implemented three genes: *APC, Kras*, and *P53*, each with two alleles, which initially start as wild-type and eventually acquire mutations based on a chosen mutation rate. Although there is the possibility of building a multi-loci model where: *A*_*n*_, *B*_*n*_, *C*_*n*_ represent the three genes in our model, we have opted to calculate the frequencies of the three genes separately. This decision is based on the absence of epistasis or recombination events in our model. After defining the genetic traits, we needed to extract not only the frequencies from the model (the number of cells bearing the phenotype divided by the total number of cells in the population), but also the fitness of each individual. In classical population genetics, fitness refers to the contribution of a given phenotype to the numeorsity of the next generation, assuming a complete correspondence between genotypic and phenotypic traits. However, the definition of the genotype-phenotype mapping can be a complex issue. It is often represented as a multi-layered graph where various genotypes form the base, with arrows pointing to one phenotype in the layer above. Importantly, the complexity arises because one phenotype can be formed by different genotypes, and one genotype can give rise to more than one phenotype, indicating a many-to-many relationship with additional parameters involved [37, 38]. In our model, the genotype-phenotype map is clearly defined. For instance, mutations in both alleles of APC yield the *pre-adenoma* phenotype, while adding a mutation in at least one allele of *KRAS* results in the *adenoma* phenotype. Finally, if both alleles of P53 are mutated, the phenotype becomes *tumoral*. As a result, the fitness values only change in two cases: the mutation of one allele of *KRAS* and the emergence of the *tumoral* phenotype. In these scenarios, the mean mitotic activity decreases to 12 hours for *KRAS* mutations and 10 hours for *tumoral* cells. Following the classical formulation mentioned earlier, we can assign the proper fitness values to every gene and allelic variant accordingly.

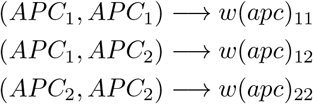

This set of relations is to be extended to the other two genes, and then putted into the recursive equation to track the allelic frequency.

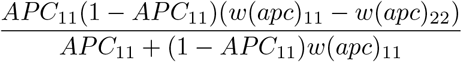

This is a slightly different version from the aforementioned equation, taken from [32], but the result is the same.

### 3.3 Results with classic selection model

In this section, we present the results obtained by setting the fitness values of the four phenotypes as random values sampled from a normal distribution. Specifically: The *normal* phenotype has a mitotic distribution time with a mean of 24 hours and a standard deviation (std) of 2. *Pre-adenoma* cells have the same mitotic distribution parameters as the *normal* cells. *Adenoma* cells exhibit a higher proliferation rate, indicating a higher fitness value. Their distribution has a mean of 12 hours. *Tumoral* cells have the maximum fitness value, with a further reduced mean mitotic time of 10 hours. With some updates ^2^, addressing issues from the previous version of the model, the settings for the runs remain consistent with those presented in our previous work:

The number of immune system cells is set at 1*/hour*. The number of cells that a killer cell can eliminate before it dies is 5. The hypoxic threshold is established at 5 *cells/patch*. The initial probability of being heterozygote is *P* = 0.5. These settings provide a foundation for analyzing the results and comparing them to previous simulations.

#### 3.3.1 Fitness embedded

In the following section, we present results obtained using the population genetic model in various scenarios. These scenarios have been kept consistent with the previous work, with all parameters remaining static except for the probability of mutation of the driver genes, which ranges from 10^*−*9^ to 10^*−*2^. Our goal is to observe not just changes in allele frequencies as the mutation probability increases—since such changes are expected by definition—but also to identify which genes are more susceptible to selective pressure. To measure the selective pressure, we have plotted the allele frequency *p* against the variation through generations of the same allele, denoted as Δ*p*. This approach enables us to observe whether there are certain allelic frequency values at which there is a positive, negative, or negligible variation in allele frequency. You can refer to Figure. 1, for a visual representation of these dynamics.

**Fig. 1.**
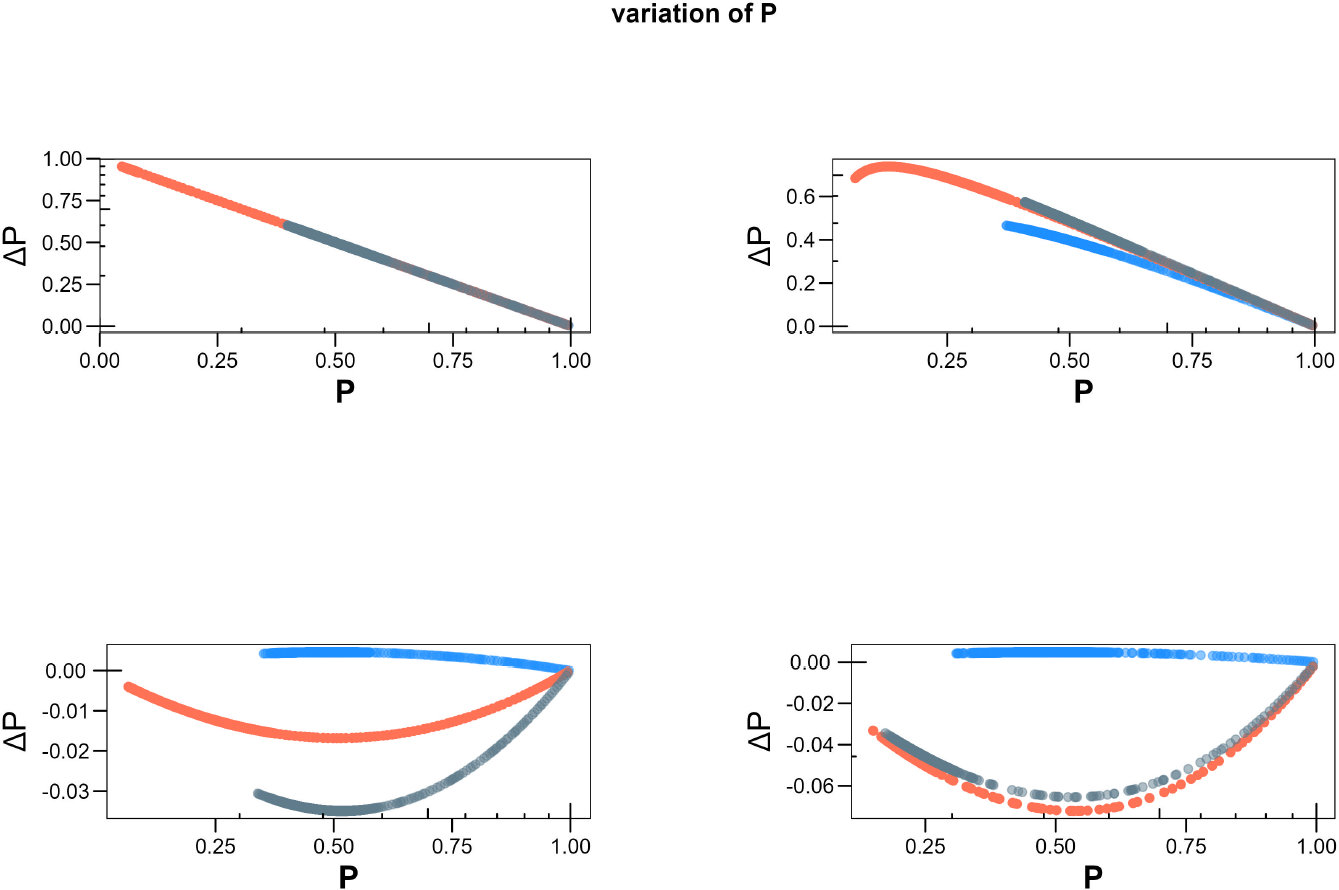
From the top left of the picture, the variation of the three wild-type alleles in the crypt at 10^*−*9^, 10^*−*6^, 10^*−*4^, 10^*−*2^ probability of mutation, in function of the frequency of the allele itself. *APC*_11_ is depicted in blue, *KRAS*_11_ in orange, and *P* 53_11_ in grey.

In the first quadrant from the left, it is easy to observe a linear relationship between Δ*p* and its frequency. Specifically, when the genotype *KRAS*_11_ is present in every cell of the crypt, its variation from one generation to the next is zero. Conversely, as its presence approaches zero, the variability becomes extremely high, indicating a population tendency towards the physiological value.

The other two genes, *APC* and *P53*, exhibit a similar trend to that of *KRAS*, differing only in the onset of variation, which occurs at a frequency of 0.5. As a comprehensive result, we can observe that Δ*p >* 0 until it reaches the maximum frequency. Therefore, the *APC*_11_ variant experiences stabilizing selection when the crypt state is physiological. Individuals with intermediate traits have a fitness advantage over those with the wild-type genotype, [39]. In the second image, both *APC*_11_ and *KRAS*_11_ begin to show variability, indicating the emergence of different phenotypes from the physiological state, particularly at a frequency of 0.4*−*5. Similar behavior is observed at 10^*−*5^, although it is not depicted in the image. Stabilizing selection persists in these scenarios, similar to what is observed in the physiological and low probability of mutation scenarios, i.e., 10^*−*8^, 10^*−*7^. The critical aspect of the trend is the initial increase in Δ*p* from 0.6 to 0.75, suggesting a change in allele frequency, which also applies to APC. This behavior might imply that, after a mild directional shift, the population eventually returns to a stable equilibrium, aligning with the characteristics of a biological system like the colonic crypt, where low mutation probabilities offer protection against harmful phenotypes, [10]. At 10^*−*4^, a pronounced curve is observed in the negative quadrant, indicating a fundamentally different behavior for physiological alleles. The variation remains in the negative quadrant and reaches zero only at a frequency of 1. This trend likely arises from the significantly reduced fitness of low trait value variants, especially between 0 and 0.5 frequency. Beyond that, there is an ascending trend reminiscent of disruptive selection, perhaps favoring two extreme phenotypes over those aided by stabilizing selection [39]. Alternatively, directional selection may be at play, altering the population’s average phenotype and reducing variation, [39].Another point to consider is the limited variability of *APC*, compared to *KRAS* and *P53*, both of which exhibit similar trends. While not explicitly clear from the data, several hypotheses can be made. For instance, the reduced variability in APC may be linked to the fitness change resulting from the mutation of *KRAS* and the role of *P53* as a cell guardian. If the cumulative effect of regulatory gene mutations reaches a certain threshold, and *P53* remains wild-type or heterozygote, it triggers cell death. Consequently, subpopulations with KRAS mutations tend to undergo more mitosis, increasing the likelihood of errors in gene lists, which leads to cell death via *P53* ‘s actions. Therefore, this variability may be a consequence of a feedback mechanism against an increased mutation probability—an adaptation. However, following a more simple explanation, we can argue that *APC* mutated alone, which is *pre-adenoma* phenotype, is less numerous in the crypt, given the fact that it does not bear any pre-imposed fitness advantage, resulting neutral to selective forces. To further support the selection model analysis, we can integrate information using population distributions from the same scenarios. The first result shows a linear trend, suggesting a preference for intermediate trait values. In the top-left picture of Figure 2, the average value shifts from 1.5 to 1.8, with a notable increase in the number of cells with that trait value. This indicates stabilizing selection with a slight increase in the average value. The second image corresponds to a scenario where the trend differs slightly for low frequencies, resulting in stabilizing selection. Although stabilizing selection typically does not alter the average value (which is influenced by directional selection), we observe the mean value increasing up to 2 for intermediate trait values, starting from an initial mean of 1.5. It’s possible that both types of selection have acted on the population, and under certain environmental conditions, less extreme values have become less fit. The third and fourth images depict crypt populations at 10^*−*4^ and 10^*−*2^, representing high probabilities of mutation. These scenarios are observed in crypts where cell DNA is affected by several epigenetic detrimental factors, such as *chromosomal instability (CIN), microsatellite instability (MSI), DNA mismatch repair deficiency (MMR)*, and *DNA methylation* [40]. The bottom-left scenario reflects negative selection against physiological traits with lower fitness. While the average value remains stable at 2, it is evident that individuals with a trait value of 0 experience a significant decrease between the two time steps. In the last image, directional selection is evident, as the initial population exhibits an almost bimodal distribution with an average of 2.5. The descendant population has an average value of almost 4, with a loss of 50,000 cells for lower value traits.

**Fig. 2.**
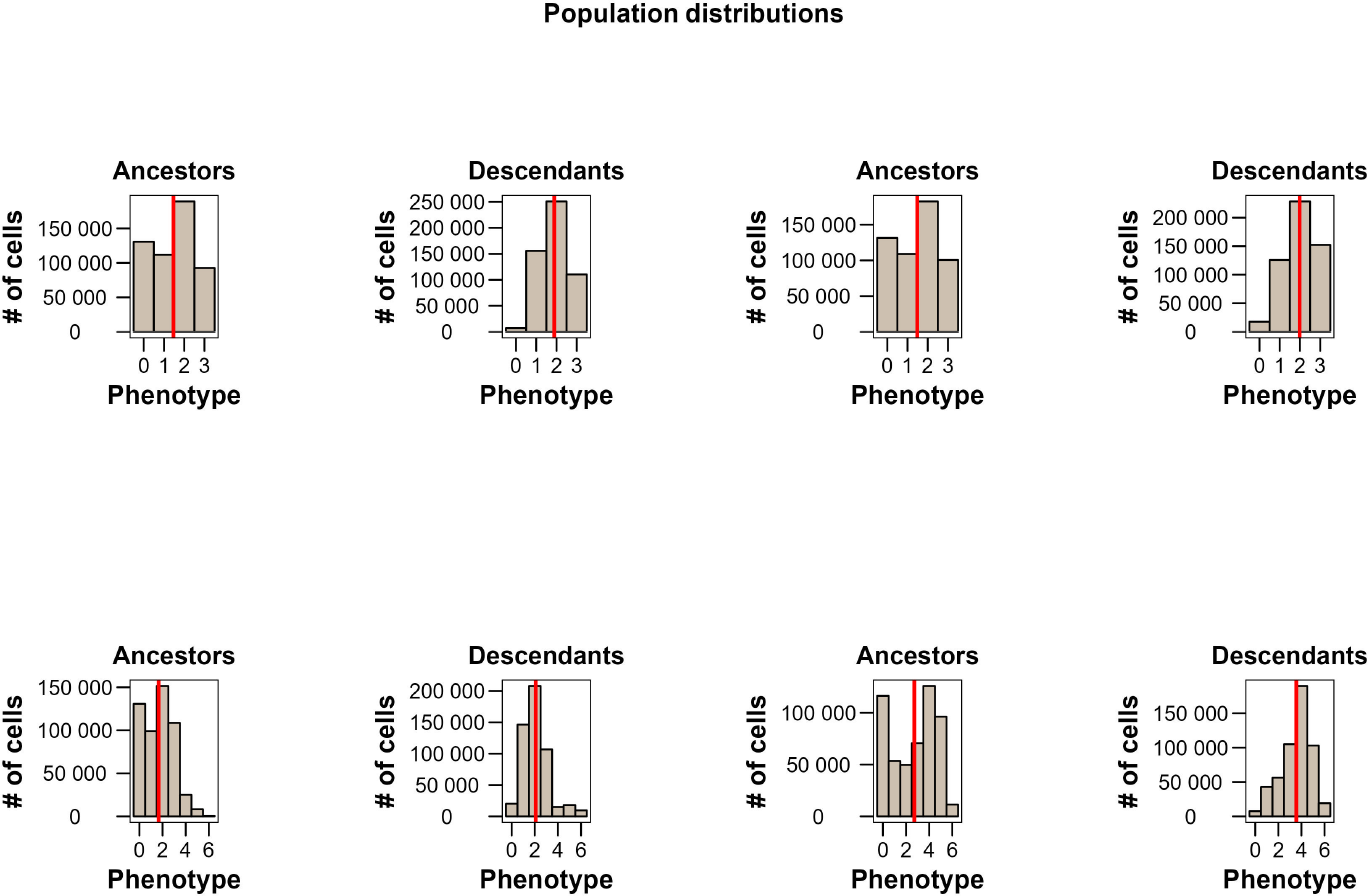
Here are represented the various distributions of the crypt population based on the probability of mutation scenarios, with fitness embedded. They are two for every scenario, divided in two time steps labeled as ancestors and descendants, in the *x* axes is reported the value correspondent to the sum of the allelic list (see the main text for details), in the *y* axes is reported the number of cells, and the red line is the average trait value of the population.

#### 3.3.2 Fitness excluded

Now we turn our attention to a scenario where we have introduced a fundamental change to the model, specifically, it has been removed the mean time of mitosis, resulting in a presumed increase in fitness, for *adenoma* and *tumoral* phenotypes. Before presenting the results, it’s essential to clarify our procedure: in the fitness-embedded model, we explicitly set the variation in the characteristic linked to proliferation, i.e., the mitotic time. However, in the current model, we have only defined the initial physiological value, with a mean of 24 hours. Consequently, the primary distinction between physiological and neoplastic phenotypes lies in the movement degrees of freedom.

With this context in mind, when we examine Fig.3, we can observe certain differences that can be attributed solely to the absence of explicitly declared fitness values when genes mutate.

**Fig. 3.**
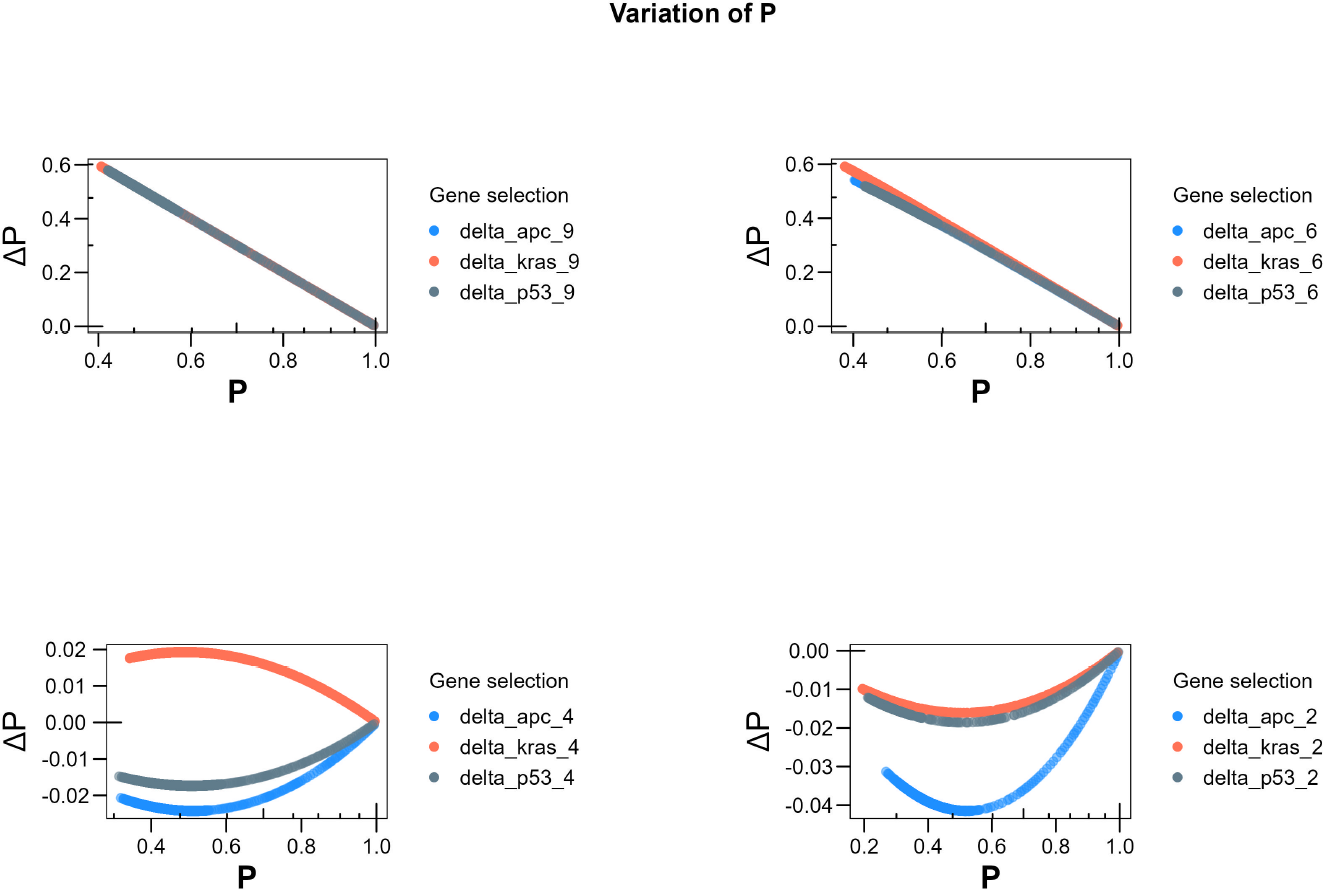
Selective trends with fitness excluded.

In the first panel, starting from the top left, we examine the scenario at 10^*−*9^. Here, we observe a familiar stabilizing tendency, akin to what we observed in the fitness-embedded simulations. An important point to highlight is that all three genotypes initiate their variability at a frequency of 0.4. This observation aligns with the absence of a fitness increase through heterozygosis in *KRAS*. Moving to the second panel, we notice a similar trend to the previous scenario at 10^*−*6^, characterized by low directionality at low frequencies. In scenarios with a higher probability of mutation, we observe significant changes in variability trends, akin to our previous simulations. Notably, *APC* and *P53* experience negative selection, escaping this selective pressure only at higher frequencies. In contrast, *KRAS*, unlike the fitness-embedded scenario, undergoes directional positive selection. This could be attributed to its ability to occupy a broader region within the crypt every time it undergoes mitosis. The trend at 10^*−*2^ roughly mirrors the pattern observed in the fitness-embedded plot, where all three genes are subject to negative selection. In this particular case, it is conceivable that cells with fitness determined by their ability to use unoccupied patches for daughter cell placement, can do so more rapidly than the rate of mitotic activity exhibited by physiological cells. From a biological standpoint, this implies that while physiological cells are stable collective entities with limited mobility and the inability to override neighboring cells without specific signaling (due to their rigid spatial structure), neoplastic cells can adopt different forms, squeezing themselves between two or more other cells, affording them greater spatial freedom.

Additionally, it’s worth noting that the population distribution for each scenario can provide valuable insights, as depicted in Fig.4.

**Fig. 4.**
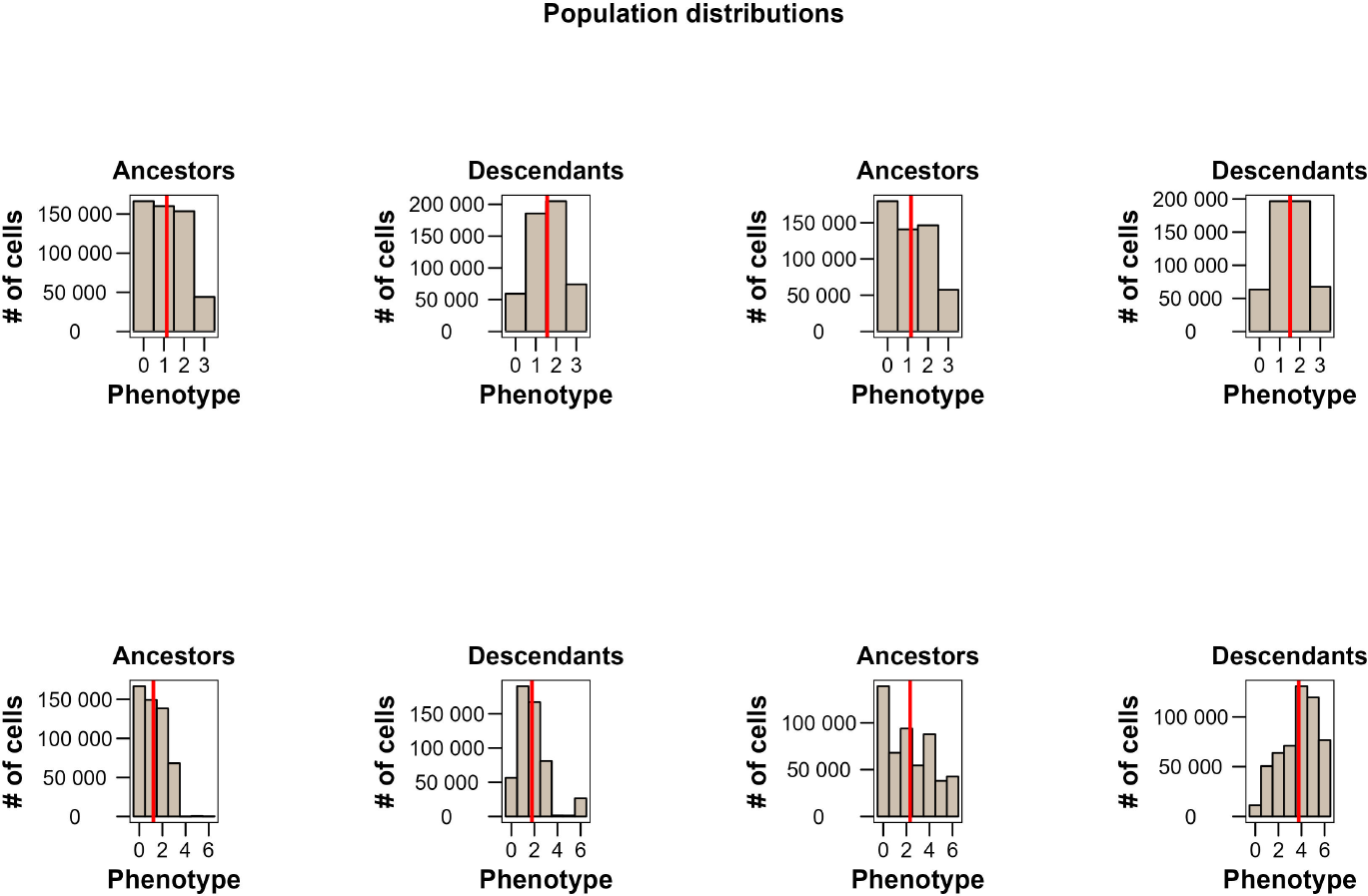
Population distributions with fitness excluded.

In the first panel, starting from the top left, we observe a stabilizing tendency towards mean trait values, with a negligible change in the average value. This trend becomes even more pronounced in the second panel, primarily because both trait values 1 and 2 are present in approximately equal proportions among the cells. In the third image, it’s challenging to pinpoint the specific reason for *KRAS* ‘s resistance to negative selection. However, what’s evident is the presence of negative selection against trait value 0. The fourth image reveals a significant variation in the average trait value, approaching nearly 4. However, there’s also a greater coexistence with other trait values, indicating a directional selection with some disruptive effects.

### 3.4 Price formalism

To uncover mutual relationships between evolving phenotypes and their relative fitness, we employ Price’s equation—a powerful tool in evolutionary biology. Price’s equation aims to reveal correlations between these variables at every level within the system under investigation. While the Fisher’s model of selection is commonly used in population genetics, Price’s equation is better suited to our model because we are not considering sexually reproducing organisms with the assumption of random mating. Price’s equation serves as a mathematical identity that holds true irrespective of its applicability. Whenever there is a hypothesis aimed at explaining the evolution or change of a biological entity’s trait, this equation serves as a valuable tool for discovering such crucial relationships [41]. Moreover, it has been demonstrated that the most abstract form of this equation is not limited to biological phenomena but extends to processes where information flows from one set to another, with the potential for errors in the replication of this information [42]. In 1970, George R. Price made a significant contribution to evolutionary biology and theoretical biology as a whole by deriving the equation that now bears his name [43, 44]. The importance of this formalization of evolutionary dynamics lies in its remarkable generality and simplicity. It is applicable to both asexual and sexual populations, to scenarios involving one or multiple alleles in different loci, and it accommodates various factors, including epistasis phenomena and considerations of group and multi-level selection, [45, 46]. Price’s equation allows us to depict the evolutionary relationships between two sets of entities that share one or more phenotypic traits and a parent-offspring linkage.

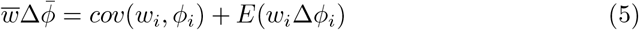

A wide literature has covered the analysis, mathematical and philosophical, of the Price’s equation ([32, 42, 47–50]) showing usefulness, [15, 51], and limits [52, 53]. In our work we have used this formalism as a machinery where put in data extracted from the model to observe if changes in gene frequencies, and increase or decrease in cell population quantities, are explainable in terms of selective pressure and importance in relative fitness. A completely exhaustive derivation of the final form of the equation is reported in the aforementioned literature, here we want to show how the equation is built and which quantities are explained by its formulation.

The 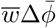 in the left side represent the mean phenotypic value scaled by mean fitness value of the population. The first term in the right side of the equation is *cov*(*w*_*i*_, *ϕ*_*i*_), which measure if and how the fitness associated with the phenotype under observation change across time. The covariance term is a very general one, capturing changes caused either by selection or drift. The second term, *E*(*w*_*i*_Δ*ϕ*_*i*_), capture the differences in trait values between parents and offspring, again scaled by fitness. The *i* indexing is the single trait (allele, genotype, phenotype etc.), or any individual identification, belonging to a particular individual of the ancestral population. Thus, if *ϕ*_1_ = *x* is the trait value in population of ancestors, 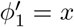 will be the sub-population in the descendant generation bearing the same trait value.

In contrast to traditional population genetics, which primarily focuses on measuring changes in allele frequencies within a population, the approach using Price’s equation is more general and can encompass a range from allelic frequency changes up to the characteristics of the entire organism within a population. In our crypt-world, the phenotypic trait we aim to track across generations is the *tipology* which can manifest as *pre-adenoma, adenoma* or *tumoral*.

In genetic analysis, two types of phenotypic values can be assigned: quantitative and qualitative. A quantitative, or metric, trait is a characteristic of an individual or group that corresponds to a measurable value, such as human height or weight. These traits, which are influenced by multiple genes, have been the subject of study since the pioneering work of Fisher and Pearson, shedding light on the clear contribution of multiple genes to the final makeup of an adult individual, [54]. However, some phenotypes exhibit discrete and qualitative characteristics, like specific durations for mitotic activity and allowable degrees of movement. To connect these properties to a quantitative aspect, we have chosen to quantify them by counting the sum of various gene lists. Consequently, the phenotypic value *ϕ* is calculated as follows:

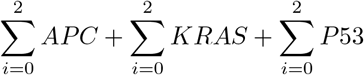

The results span from *ϕ* = 0, representing the physiological trait value, to *ϕ* = 6, indicating the *tumoral* trait value. Between these two extreme values lie various possible combinations, some neutral and others corresponding to neoplastic variants. For instance, when 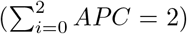, it corresponds to the *pre-adenoma* phenotype. When 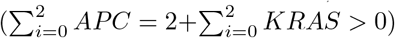, it corresponds to the *adenoma* phenotype. Finally, when 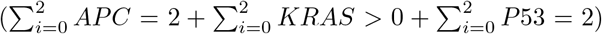, it corresponds to the *tumoral* phenotype. All other combinations are neutral regarding changes in movement. However, there is a combination involving only *KRAS* that increases the mean mitotic time, i.e., 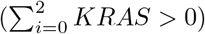.

The linkage between changes in phenotype and subsequent fitness changes is expected in this model, as it reflects how we have programmed the cells to behave. Nevertheless, valuable insights can be gained by applying this analysis specifically to the *adenoma* and *tumoral* phenotypes. To perform an analysis of the model using the Price equation, we need to divide the total population into two sub-populations based on an arbitrarily chosen time interval. It’s worth noting that this procedure may not capture all possible variations between individuals within and outside of this time interval. One can choose different time intervals, either longer or shorter, to examine evolutionary differences between the two sets of populations. In the model, agents need to be divided into these two sub-populations. To prepare the dataset, each cell is assigned an identification number, *i* = *randomid*(1, …, 5000), belonging to the single cell, and its trait value, *ϕ*_*i*_ = (1, …, 6), which is inherited by all daughter cells unless mutations occur during mitosis. The results of this analysis are depicted in Fig. 5.

**Fig. 5.**
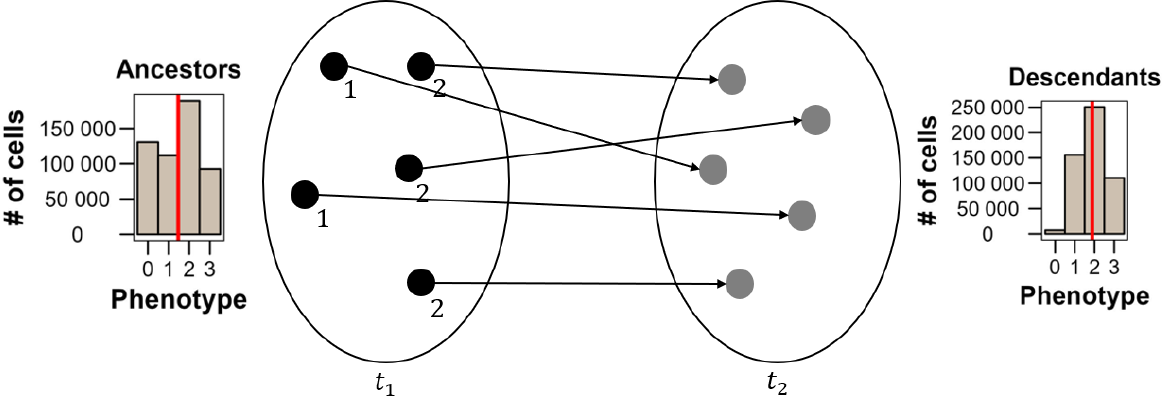
Here is depicted the relation of ancestor-descendant divided into two sub-populations. The individuals at *t*_1_ are the *ancestors* while the ones at *t*_2_ are the *descendants*. The label, which is carried by each individual, represent the phenotype value and the arrows are pointing to the belonging descendant of each of them.

We now have two populations at different points in time on which perform the necessary operations. The procedure begins by calculating the covariance between the trait values of the entire population of *ancestors* and their corresponding fitness values. In accordance with the classical definition in evolutionary biology, fitness is defined as the number of offspring in the next generation. The subsequent step involves determining the trait difference between the number of offspring in the descendant generation and the number of offspring in the ancestral generation of cells. Hence, the Price equation is rewritten in the form

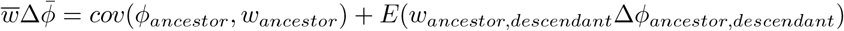

Given that daughters cells of an individual *x*_*i*_, will have the same identification number inherited from the parent, we can plot the population trait value in the future moment, namely the descendant sub-population, on the ancestor sub-population, as in the classical approach of dynamical systems, where the system state at *t* is the result of an iteration of the system at time *t−*1. Price equation gives useful information about the evolution of trait values as shown in Fig.6, where is plotted the change between generations in the scenario with probability of mutation set at 10^*−*4^.

**Fig. 6.**
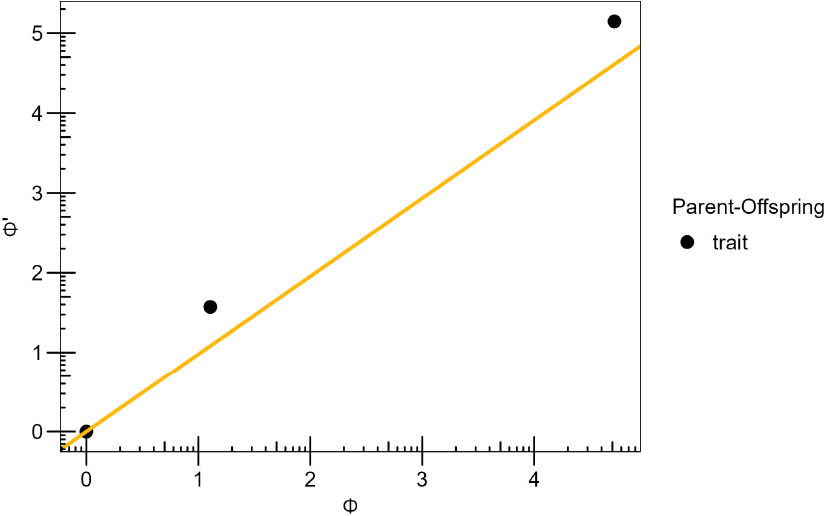
The plot shows three data points representing the minimum, average and maximum trait values of the two sub-populations. The straight line is a regression line, with intercept the minimum trail value, and slope mean change in trait value, which is the result of the Price equation.

It is crucial to emphasize that the first term, *cov*(*ϕ*_*ancestor*_, *w*_*ancestor*_), measures a statistical association, accounting for the change directly attributed to differences in the contribution of the ancestor population to the size of the descendant population, in other words, how strong is the selective effect is on that trait. In contrast, the second term, represented as *E*(*w*_*ancestor,descendant*_Δ*ϕ*_*ancestor,descendant*_), signifies the alteration in the resemblance between ancestors and descendants, which can arise from various events, both within and outside individuals. What we expect is the discovery of higher covariance values as the probability of mutation, and consequently, the likelihood of neoplastic phenotype occurrences, increases.

#### 3.4.1 Fitness embedded

Results are obtained changing the probability of mutation of the genes - *APC, KRAS, P53* -, and extrapolating data about the fitness of the phenotypes in function of their trait values, as depicted in Fig. 7.

**Fig. 7.**
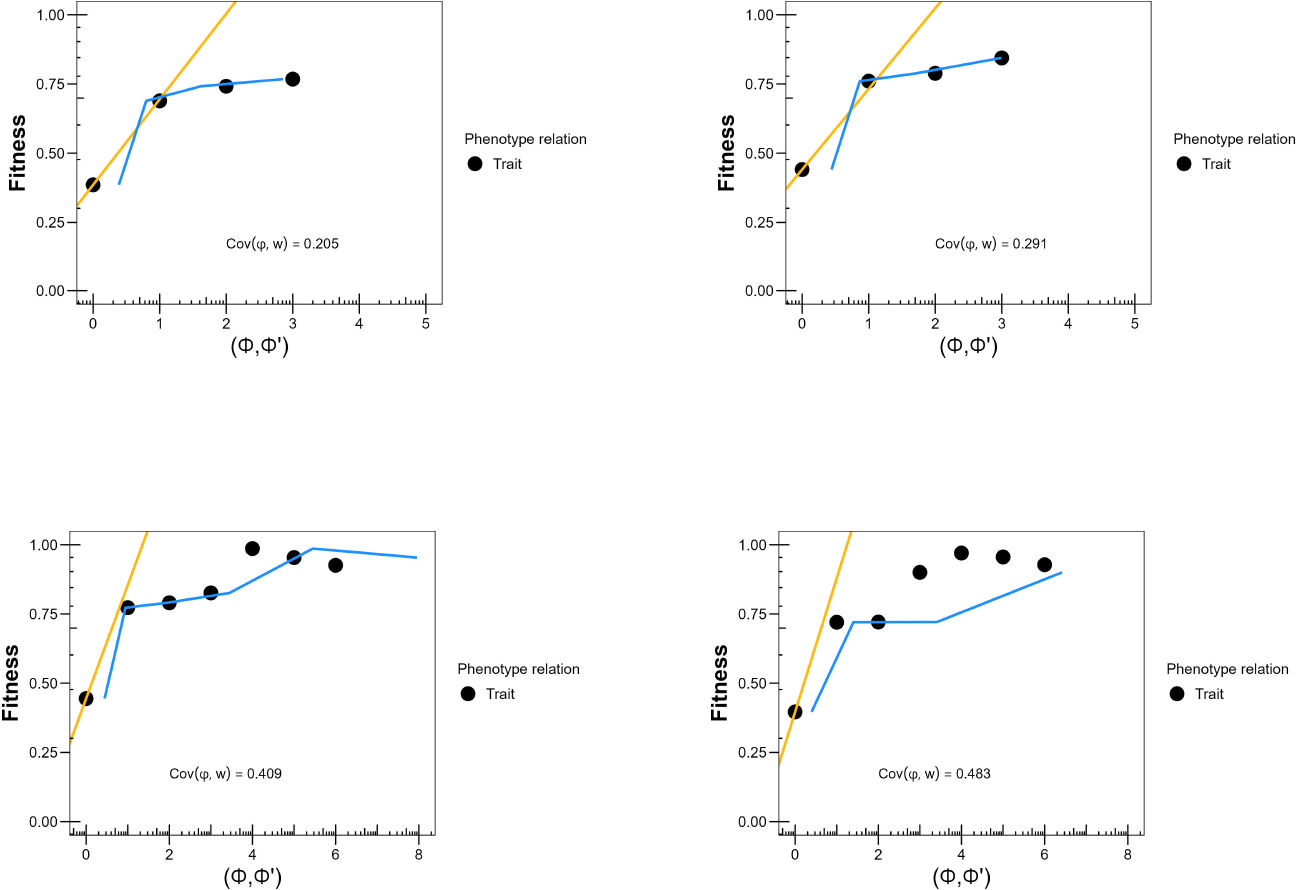
Scenarios are the same as for the classic genetic model, probabilities of mutation 10^*−*9,6,4,2^. In the *x* axes are reported the trait values, in *y* axes the fitness scaled on the maximum fitness of the entire population. The golden line represent a linear regression line the azure one, is obtained with a second order polynomial to seek the parabolic trend, see the main text for details.

**Fig. 8.**
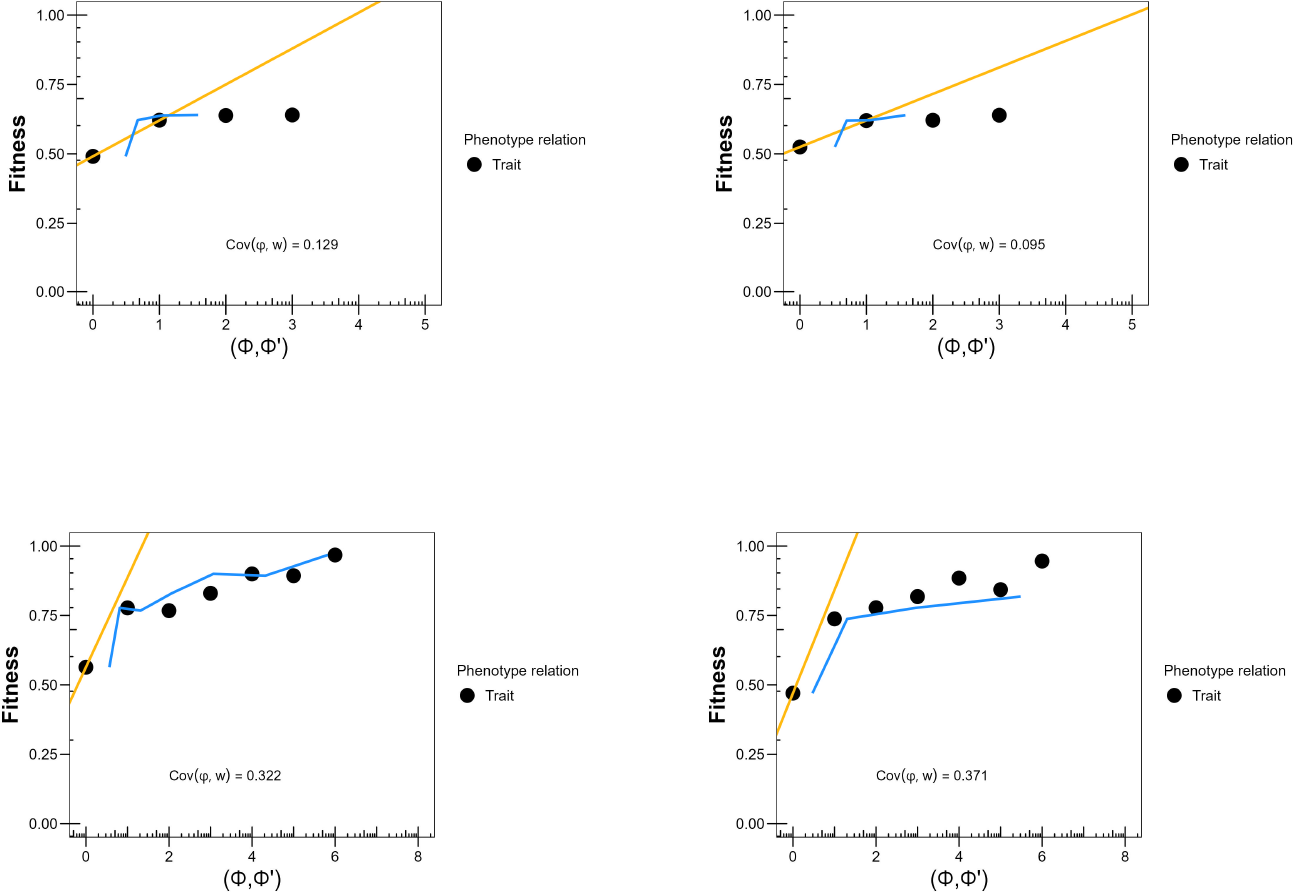
Scenarios are the same as for the classic genetic model, probabilities of mutation 10^*−*9,6,4,2^. In the *x* axes are reported the trait values, in *y* axes the fitness scaled on the maximum fitness of the entire population. The golden line represent a linear regression line the azure one, is obtained with a second order polynomial to seek the parabolic trend, see the main text for details.

**Fig. 9.**
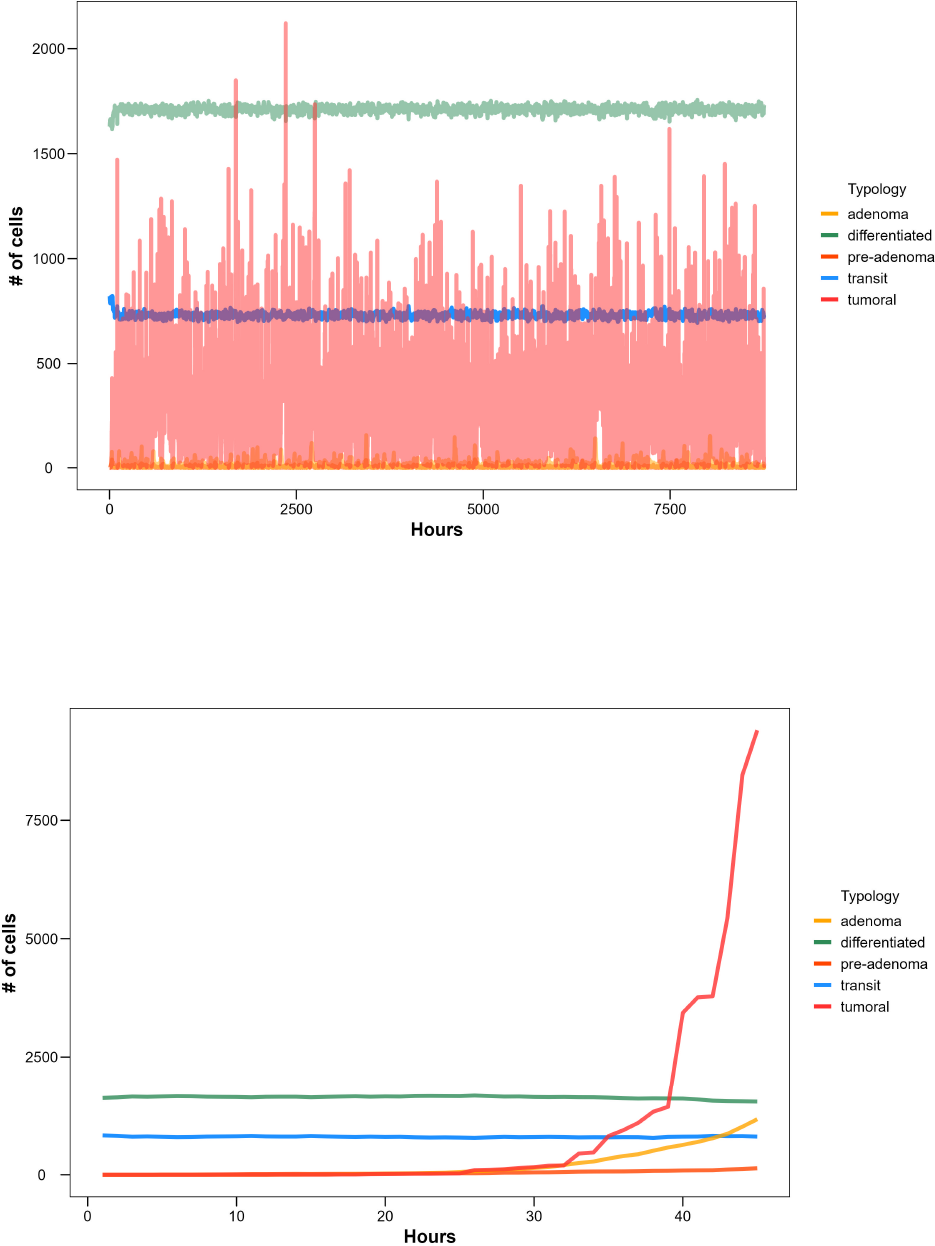
Crypt at 10^*−*2^ probability of mutation, with and without environmental constraints.

Now, let’s delve into the various scenarios and the biological implications they represent. As demonstrated in different studies, the *β* term of the equation pertains to changes in the selection of the trait. This term reflects alterations in fitness values that occur as an effect of changes in the trait value [32, 50]. This information is clearly presented in the plot, where we observe a notable increase in covariance values as the probability of mutation rises, ranging from around 0.3 in the physiological scenario to nearly 0.5 in the completely deregulated crypt at 10^*−*2^. The fact that the covariance term is not 0 in the physiological crypt is probably caused by the probability of heterozygosis, in the bottom line cells, set up at 0.5, meaning that each of the 25 original cells has a good probability to be heterozygote for each one of the three genes, eventually reaching *ϕ* = 3 even from the very beginning of the simulation. The lower covariance at 10^*−*6^ probability of mutation can be explained by the rarity of genotypic trait values greater than 3, associated with the specific *adenoma* phenotype, at this low mutation rate. Furthermore, the increase in fitness aligns with the attainment of the maximum trait value. In contrast to the first scenario, where fitness linearly grows with the trait value from 1 to 3, once the genotype with heterozygous *KRAS* is fixed in the population, the maximum fitness remains constant along the trait value axis. In the third and fourth images, we observe a significant jump in fitness value up to 1, corresponding to the trait value 4 followed by a slight regression to 0.8 This phenomenon occurs because, at *ϕ* = 4 the more likely phenotype is *adenoma* which contributes to altered movement and increased mitotic activity in the cells. The reduction in fitness values for *ϕ >* 4 can be explained by the characteristics of the *adenoma* and *tumoral* phenotypes encountering environmental elements, including the hypoxic threshold and immune system cells. When the *adenoma* phenotype arises, cells exhibit a random 90-degree movement randomly directed to the left or to the right. This altered movement reduces the time it takes for these cells to reach the crypt’s mouth. However, it’s not low enough to make them easily preyed by killer cells, or for allowing them to accumulate in patches and trigger the hypoxic threshold. This ability to evade environmental threats is crucial for maintaining a higher number of descendants. On the other hand, *tumoral* cells have a longer average mitotic time but no longer migrate outside the crypt. Instead, they proliferate in nearby patches with a 360-degree degree of freedom. Unfortunately for them, this extensive proliferation triggers environmental conditions that reduce *tumoral* cell numbers. Once located, these cells cannot escape killer cells, and overcrowding becomes an issue.

#### 3.4.2 Fitness excluded

Now we focus our attention to the simulations without explicit fitness values associated with phenotypes. The results here exposed shows the relevance of the covariance term to make reliable prediction about the association among trait values and fitness values, see Fig.3.

The first two scenarios, in the top row, report very clear information about the importance of altered movement to augment the fitness value, in fact, while the physiological fitness is stable at 0.5, when the trait value become *ϕ* = 1 we observe a mild increase in the fitness. Since *ϕ* = 1 can be associated only to one genotype with a different behavior in respect of the physiological one, i.e., 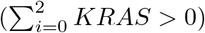, we can conclude that the possibility to find new niches among, or above, normal cells without respecting space boundaries, is an advantage, hence, a factor of fitness variation within the population. In the other two we assist to a trend resembling the one of the fitness embedded simulations. However, the main difference is in how the fitness values increase, with a more linear trend consistent with the no more present discrete assignment of fitness value. The first leap from the physiological fitness goes to 0.75 for *ϕ* = 1, then continues going up till *ϕ* = 6 where we observe the maximum fitness value of 0.9/1. Remarkably, the highest fitness value does not corresponds to the highest in the previous simulations. How is that possible? In the fitness embedded simulations, *tumoral* phenotype has a mean mitotic time of 10 hours, while now, maintains only the altered nearby movement, without caring about the location or the number of other cells, either physiological or neoplastic.

One possibility could be that, thanks to the lack of a preimposed fitness advantage, *tumoral* cells enter in mitotic activity only when their cycle is complete, every 24 hours, but, unlike the physiological cells, they do not need to use other procedures (cells are allowed to proliferate only towards free patches, if they are occupied and the mitotic time is activated, they choose one of the neighbors and kill it based on neighbor’s fitness or age), to fill the adjacent patch. Therefore, they can proliferate more than both, physiological and *adenoma* cells, overtaking the latter whose movement is more directly oriented towards the crypt mouth without enough time to proliferate with out the aid of an artificially augmented fitness.

### 3.5 Insights on results

The significance of discovering general relationships between trait values and fitness, beyond those already established in the model, led us to develop some mathematical treatments of the data. In the plot, two regression lines are depicted to fit the data points: one is a linear regression built with the minimum fitness value between the two sub-populations as intercept, and the *cov*(*ϕ*_*ancestor*_, *w*_*ancestor*_) from equation (4) as slope; the other is a second-order polynomial designed to capture the curved trend. Both linear and non-linear regression lines were developed with the aim of identifying parameters with generality within the model, seeking invariant quantities with predictive value [55]. The most fitting function we found combines the covariance value of the parent sub-population, using it as a second-order term and as a first-order term, with the minimum fitness value between the two sub-populations as the intercept.

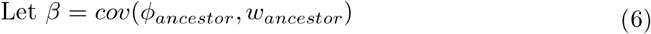

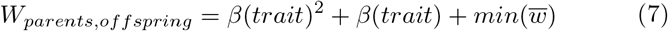

Now, the challenge lies in attributing biological significance to this mathematical formulation, and achieving this would be a valuable accomplishment since the aim of these analyses is to uncover meaningful evolutionary causal relationships between trait and fitness values. The purpose to find such solution to this issue, is to view the function as a predictor, stemming from an ancestral generation, of the fitness values for a future generation. This perspective allows us to understand how changes in trait values impact the increase or decrease in the contribution of offspring. From a dynamic standpoint, this implies that we can begin with an initial state characterized by the minimum fitness value associated with a particular phenotype. As time progresses, the changes in the phenotype, along with the corresponding fitness, as captured by *β*, result in the subsequent state of the population. This approach allows us to forecast how the population’s traits and fitness values may evolve over time, shedding light on the evolutionary dynamics within the system.

## Discussion

Considerable attention is drawn to the concept of natural selection and its significance in an artificial environment. As Charles Darwin himself pointed out, the principle of selection comes into play when there is the transmission of information and differences in that transmission. It resembles the action of artificial selection carried out by breeders on their animals. However, in nature, this process occurs as a struggle for survival and reproduction among various living organisms, as described in [56]. Therefore, it is a non-randomized process that directly influences the evolution, i.e., the change in one or multiple characteristics from one generation to another, of the elements involved in the process. From this perspective, one might legitimately ask, what are the constituent elements of this process? The most reasonable answer is a combination of heritable and ecological factors whose action results in limiting population growth. If any biological population were left to proliferate unchecked, it would exhibit exponential, geometric growth, which would be unsustainable for the environment in the absence of limiting factors. For example, consider the bacterium E. coli, capable of reproducing every 30 minutes. Given infinite resources, it could produce an astonishing number of copies of itself. Similarly, as noted by Darwin, even a single elephant could give rise to nearly 20 million descendants in just 750 years, as described in [57]. This exponential growth is also observed in cellular populations, such as those eventually reaching a neoplastic phenotype, where the overall functional behavior of the tissue is no longer solely the result of individual cell actions. Early hypotheses about tumor progression considered tumoral cells as proliferating without constraints. However, it has been demonstrated that this exponential growth pattern only holds for the initial part of population increase. In reality, biological populations are limited by a factor known as the carrying capacity of the system, as outlined in [58].

Such upper level interactions are described, in the present model, as population constraints, and predator-prey relations [59], responsible, as well as probability of mutation, for the selective pressure on certain phenotypes from others. As observed in both classical genetic model and Price equation manipulations, where there is a considerable presence of inter-specific interaction and death of cells due to overcrowding, we observe selective pressure in all the three genes involved, in both simulations with and without fitness assigned values.

Let’s consider an example from the present model. When we observe population dynamics in the presence of environmental conditions, such as immune system cells hunting neoplastic cells and a hypoxic threshold causing the death of basal cells, we notice a tendency toward a homeostatic state, characterized by a struggle between cellular populations (see Fig.9).

Conversely, when we remove the triggering of immune system cells and increase the hypoxic threshold to thousands of cells, the population of tumoral cells explodes in just a few hours. Therefore, we can conclude that these elements align with the thesis that cells can be considered as individuals of a particular species subject to internal and environmental constraints, and consequently, to natural selection. If there are more cellular phenotypes when immune system cells are active, then we can think about some kind of *force* acting on the individuals, causing their actual distribution. Even though it would be possible, in principle, to program cells to act with a feedback-driven behavior producing a sort of higher level self-organization, this is not the case. Cells are not programmed to react as agents, hence, every top-down constraint, or channel, is a purely blind process sprouting from the population down to the individual level.

Notwithstanding conceptual issues about the concept of *force*, when applied to natural selection and evolutionary process [60], there is room for an account of natural selection as a physical mechanism and not just the fact that something is happened to the trait distribution. As showed by [61], the selection forces against and for a given trait, can be considered vectorially, in fact, when no forces are acting on the trait, genetically and environmentally, there is no change in the trait dynamic. One significant challenge arises when attempting to define the entities under selection, a problem that has puzzled biologists since Darwin’s time, as discussed in [47]. In the hierarchical view of biological systems, entities are not isolated; they are nested within collectives exhibiting structures and functions distinct from those of their constituent units. Cancer is typically considered a multi-level phenomenon, spanning from the structures of cellular nuclei to the tumor microenvironment, which still lacks a clear spatial and relational definition, as noted in [62]. In this view, the roles of transit-amplifying and differentiated cells are to maintain the crypt population in equilibrium, secrete mucus, and absorb nutrients from substances flowing down the intestine. The function of the entire crypt and the entire colon is to ensure various operations resulting from the coordinated behavior of multiple cellular types [63] and even bacterial populations.

In other words, the physiological architecture of the crypt is co-constructed by multiple biological agents to perform functions beyond the individual agents’ behaviors. When we discuss the fitness value of individual cells in this context, we refer to the number of offspring allowed by the specific genotype and the constraints imposed by the population and microenvironment. This is why when cells acquire somatic mutations that alter this behavior, they are referred to as “cheater” or “renegade” cells, because the selective process no longer benefits the entire tissue; instead, it advantages the single lawless cell. For these neoplastic variants, collective behavior no longer serves as a fitness enhancer, as explained in [64].

Another feature worth a little examination is the selection pressure of wild-type alleles. It is known that genotypes with beneficial traits are selected out within the population, increasing their frequency and, perhaps, eventually substituting the original genotypes. In our model there are two types of selective pressures: probability of mutation and environmental elements. The first one has less importance in its change for the reason that each scenario has a stable probability of mutation, implying that cells produced in the crypt base will always bear the starting probability of mutation, not greater nor lower. However, when the probability of mutation is sufficiently high,mutation in *P53* can trigger the incremental increase in that probability: for example, if in the scenario at 10^*−*6^ a mutation on both *P53* alleles occurs, the bearing cell would have daughter cells with a probability of mutation increased by some *x*, which in our simulations is set up at *−*0.5. Such events are responsible of change in allele, genotype or phenotype frequencies in the measure that this particular occurrence is diffused in a relevant fraction of the entire population. In real biological systems, physiological states resist at both individual and population cellular level, to events who could increase probability of DNA damage, [65], to ensure the correct accomplishment of the various functions. Hence, we can conclude that, sometimes, the increased probability of mutation, in the model, is not the result of an external manipulation, is the effect of a gatekeeper gene on the probability of mutation of the others. These features, even if extremely simplified and incomplete, mirror the complex interplay of factors acting from DNA level, probably even sub-molecular level, and upper levels where inter-specific interactions gain a huge relevance.

## 4 Conclusions

In this study, we conducted several simulations with the objective of gaining valuable insights into the presence of natural selection mechanisms and how these mechanisms impact the behavior of both individual cells and the cellular population. The results appeared to indicate that environmental constraints play a pivotal role in the intricate interplay among various neoplastic phenotypes. These constraints are notably absent in crypt simulations where immune system cells and hypoxic thresholds are inactive. Hence, we can deduce that when considering an evolutionary perspective of tumoral progression, we must regard environmental factors not merely as boundary conditions but as genuine causal factors akin to the activation of oncogenes and the silencing of tumor suppressor genes. Our future research endeavors are geared towards a more detailed exploration of cellular-environment interactions, encompassing aspects such as energy consumption and cellular functions, including the secretions of differentiated cells. This approach aims to construct a structural and functional architecture whose behavior can be analyzed using multilevel selection methods.

## Appendix A Supplementary information

Model, dataset and code used in the present work are available online: https://doi.org/10.5281/zenodo.10035320

## Footnotes

1 Aristotle specifically spoke about, material, formal, efficient and final causes. The bricks of which a house is composed, the project of the house, the builder, the purpose to build a house.

2 In the previous work’s model [8], there were some variations in the cell count and the movement of adenoma cells. Regarding the first difference, we decided to modify the cell count due to an unintended consequence of the model. This unintended consequence was the accumulation of transit-amplifying cells over the stem cells. This accumulation occurred because there were no available free patches ahead where cells could move. Consequently, the cells entered a sort of stagnant state until a free patch became available for them to occupy. As for the movement of adenoma cells, we opted to increase the radius of their inclined movement. This adjustment was made to simulate a stronger attachment to the tissue and, as a result, slower movement of these cells.

